# Colourfulness as a possible measure of object proximity in the larval zebrafish brain

**DOI:** 10.1101/2020.12.03.410241

**Authors:** Philipp Bartel, Filip K Janiak, Daniel Osorio, Tom Baden

## Abstract

The encoding of light increments and decrements by separate On- and Off- systems is a fundamental ingredient of vision, which supports the detection of edges in space and time and makes efficient use of limited dynamic range of visual neurons [1]. Theory predicts that the neural representation of On- and Off-signals should be approximately balanced, including across an animals’ full visible spectrum. Here we find that larval zebrafish violate this textbook expectation: in the fish brain, UV-stimulation near exclusively gives On-responses, blue/green-stimulation mostly Off-responses, and red-light alone elicits approximately balanced On- and Off-responses (see also [2–4]). We link these findings to zebrafish visual ecology, and suggest that the observed spectral tuning boosts the encoding of object “colourfulness”, which correlates with object proximity in their underwater world [5].

---

To begin, we measured high-acuity spectral sensitivities of larval zebrafish brain neurons by two-photon imaging, capturing n = 11,967 Regions Of Interest (ROIs) across the brains of n = 13 6-7 days post fertilization zebrafish (elavl3:H2B-GCaMP6f, Figure 1A, Figure S1A-C). To record the entire brain along its natural 3D curvature we used a non-telecentric mesoscale approach coupled with “intelligent plane bending” enabled by rapid remote focussing (Supplemental Video S1, Figure S1A) [6]. A custom hyperspectral stimulator consisting of 13 spectrally distinct LEDs opposing a diffraction grating and collimator for collection [7] allowed wide-field stimulation, which was approximately aligned with one eye’s retinal acute zone. Regions of interest corresponding to individual and/or small groups of similarly-responding neuronal somata were extracted from each recording, then quality filtered, denoised and decomposed into On- and Off- responses (Figure S1A-G, Supplemental Experimental Procedures).

**Figure 1.**
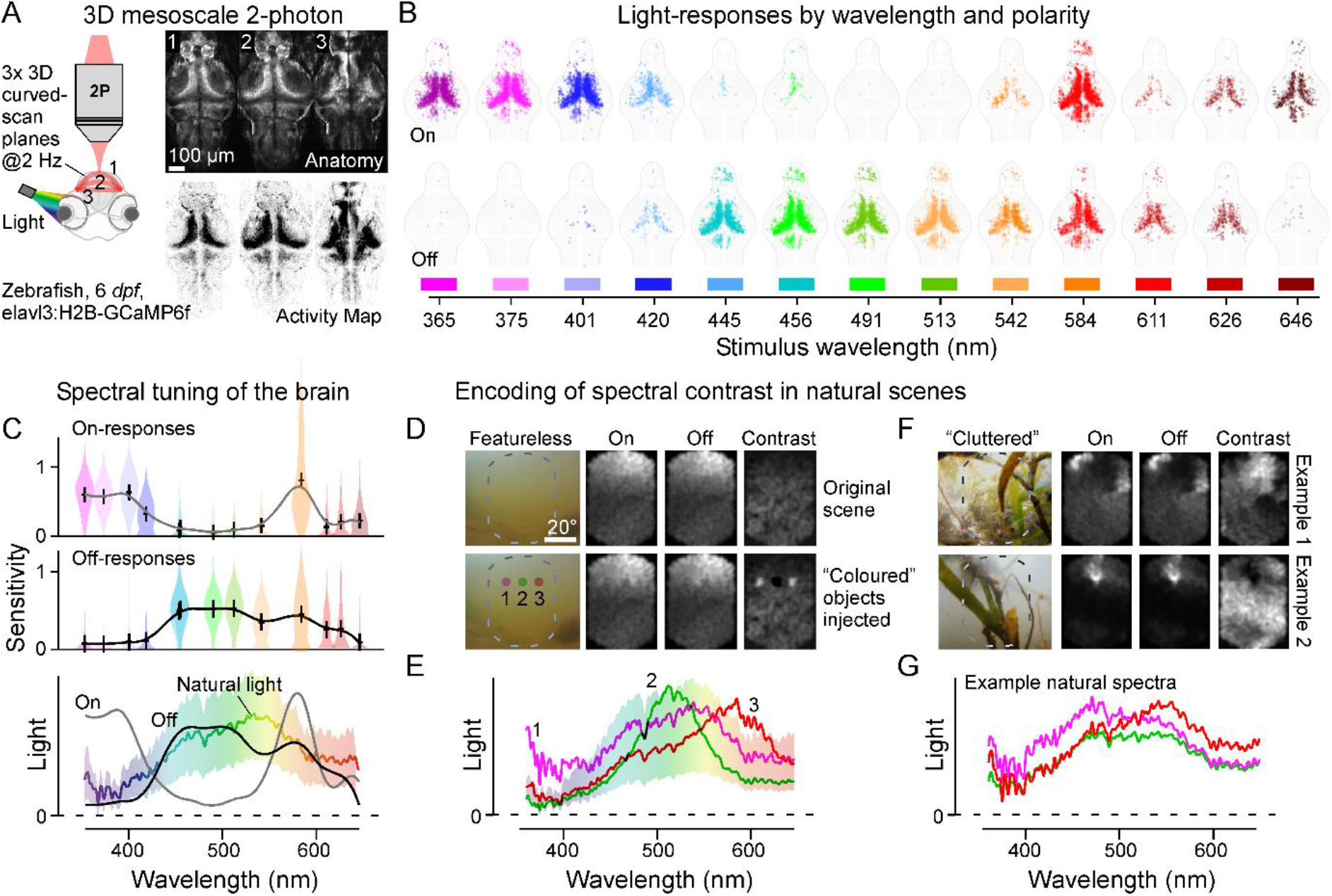
**A,** Left, larval zebrafish expressing GCaMP6f in neuronal somata were imaged on a custom volumetric mesoscale two-photon system with 3D multi-plane-bending to follow the brain’s natural curvature (described in [6]). Visual stimulation was 3 s flashes of widefield light in 13 spectral bands (described in [10]). An example brain wide quasi-simultaneously acquired tri-plane scan average (right, top) is shown alongside a projection of pixel-wise activity-correlation (right, bottom; dark: higher correlation). See also Figure S1. **B**, x-y superposition of all On- and Off-responsive ROIs (top and bottom, respectively) across n = 90 planes from n = 13 fish to flashes of light at the indicated wavelengths. **C**, Mean On- and Off-tuning functions based on (B), with crosses showing the median, and violin plots summarising the spread in the data at each wavelength (top, middle), and both tuning functions superimposed on the mean±SD availability of light in the zebrafish natural habitat (data from [8]). **D-G**, selected natural visual scenes from [8], in each case showing an indicative photograph of the scene, followed by the full hyperspectral image as seen through the On-, Off- and On-Off-contrast filters (D,F) and associated full spectra (E,G). The bottom panels of D are identical to the top with the addition of artificially “injected” local spectral distortions as indicated in E to mimic, from left to right, a “UV-“, “green-“, and “red-object”.

Recordings revealed that, despite some expected variation (e.g. Figure S1B, [2–4]), neural responses in all major visual centres of the brain had a common, overarching spectral sensitivity profile: UV-On, Blue/Green Off, Red On-Off (Figure 1B). This organisation into three spectral processing zones (UV, Blue/Green, Red) can be linked to zebrafish visual ecology. First, in the UV the dominance of On- responses likely serves prey-capture, as aquatic microorganisms appear as UV-bright objects when illuminated by the sun [7]. Second, the approximate balance of red On- and Off-responses may allow zebrafish to use the abundance of long-wavelength illumination in shallow water [8] to drive “general-purpose” achromatic vision, including motion circuits [9]. Third, the dominance of Off responses to blue and green wavelengths may serve as a subtraction signal to spectrally delineate the red- and UV-systems [2], and also to provide a spectral opponent signal for colour vision against UV- and red-On circuits [10].

A further non-mutually exclusive interpretation is that spectral organization in zebrafish brain acts to accentuate “colourfulness”, possibly as a cue to object proximity. This is because unlike air, turbidity in aquatic environments rapidly attenuates both achromatic and chromatic contrasts with distance [5], so that any high-contrast and/or colourful underwater object must be nearby.

To explore this idea, we computed the mean zebrafish brain On- and Off- spectral sensitivities and compared them to the average availability of light in the zebrafish natural habitat from [8] (Figure 1C). This revealed a good match between natural spectra and the brain’s Off-filter, while the On-filter instead peaked beyond the range of highest light availability. Nevertheless, the generally positive rectification of brain responses (Figure S1D,E,G) meant that the sensitivity of both the Off- and the On-filter strongly correlated with brightness (Figure S1J,K). Accordingly, either filter in isolation encoded achromatic infromation, which dominates natural scenes. This correlation however also meant that when computing On-Off contrast (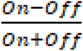) as a function of wavelength, brightness information was essentially cancelled to instead highlight spectra that differed from the mean – i.e. chromatic information (Figure S1L).

To illustrate how such an On-Off contrast filter would serve to highlight “colurfulness” in nature, we reconstructed individual natural scenes from hyperspectral images. In each case we computed three reconstructions: On-filter alone, Off-filter alone, and On-Off contrast (Figure 1D-G). In a featureless scene along the open water horizon, both the On- and Off- reconstructions were dominated by the vertical brightness gradient, while the On-Off reconstruction showed approximately homogeneous activation (Figure 1D, top). We then artificially skewed the underlying spectra of three neigbouring regions in the same image to minic small UV-, green- and red-biased objects, respectively, and again computed the On-, Off- and On-Off representations (Figure 1D, bottom, cf. Figure 1E). This manipulation had only minor effects on the On- or Off-reconstructions, however the contrast reconstrauction now readily reported the presence of all three objects. Similarly, On-Off contrast reconstructions also lended themselves to reporting foliage in the foreground in non-manipulated, cluttered natural visual environments (Figure 1F,G).

Taken together, our data suggests that the zebrafish brain’s overall spectral On-Off tuning may be well suited to compute the presence of spectral information that differs from the mean, and thus provide a useful cue to object “colourfulness”, which in turn corrleates with object proximity [5]. Notwithstanding, beyond this overarching spectral response profile, we may expect substantial additional spectral diversity at the cellular and neurite level to support the zebrafish’s diverse visual requirements [2–4].

---

**Supplemental Figure S1.**
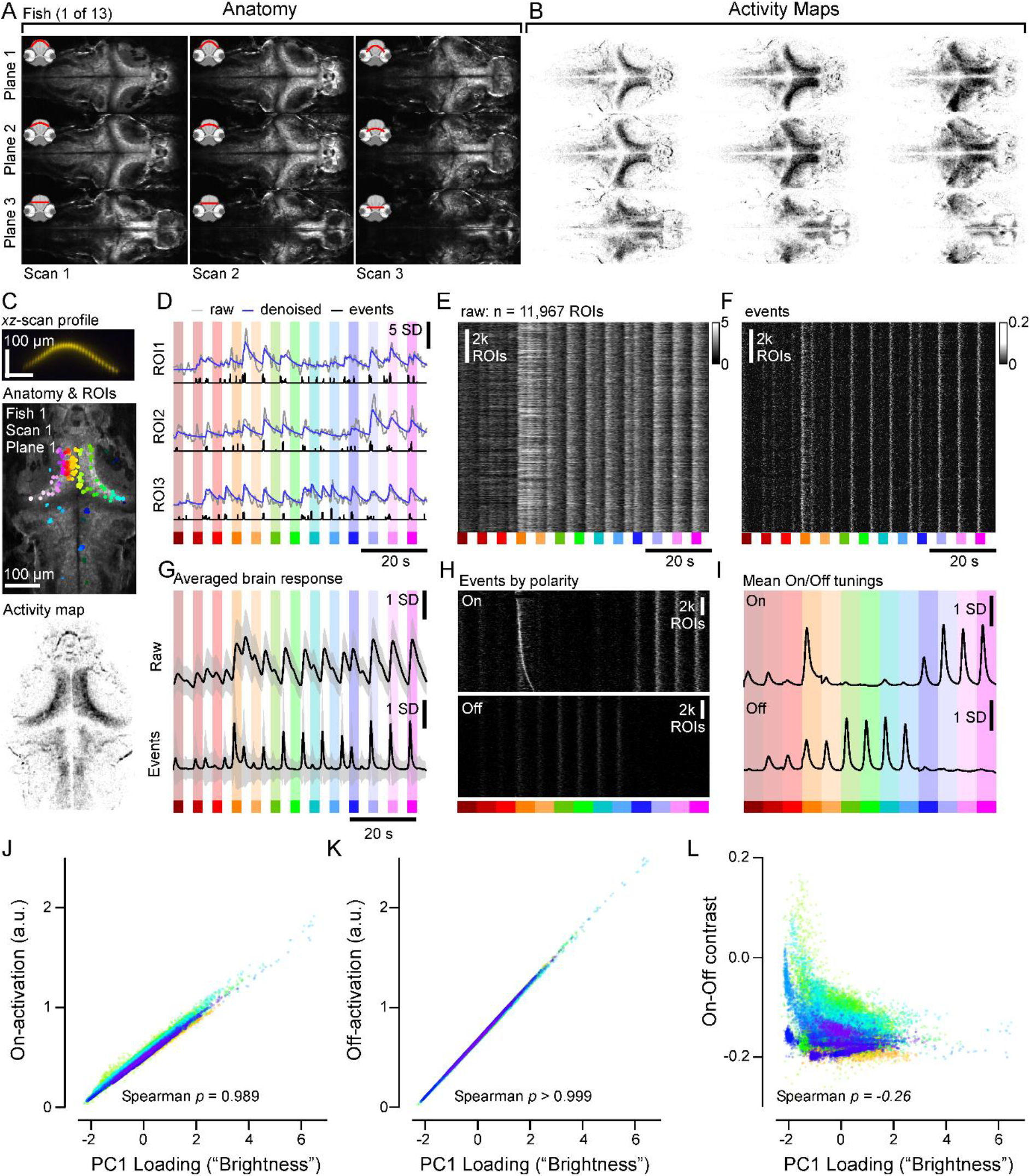
**A,** Example recordings from one larval zebrafish, comprising three consecutive scans of three planes each for a total of nine planes. For each triplane scan, starting from a common z-position, the first two planes were bent upwards by ~100 μm and ~50 μm at the apex, respectively. The lowermost plane was kept flat. Between scans, the entire triplane was moved down by ~50 μm. In total, we recorded from n = 13 fish in such a configuration. **B**, Pixel-wise activity-correlation over time with the four neighbours computed as in [S7] as an indication of locally correlated activity in the scan (darker shade indicate higher correlation). **C**, Example ROI extraction shown for plane/scan/fish 1 (cf. A,B), with xz-scan-profile visualised as in [S3] (top), a crop of the anatomical projection with ROIs (middle) and corresponding activity map (bottom). **D**, Example ROIs from (C) in response to light-flashes of different wavelength as indicated, shown as z-normalised fluorescence (grey), denoised (blue) and detected events (black). **E,F**, All n = 11,967 ROIs from 13 fish (30 scans) shown as raw fluorescence (E) and as events (F). Note polarity switches between light-flashes of different wavelengths. **G**, Mean±1SD z-normalised fluorescence (top) and events (bottom) of all ROIs. **H,I**, Separate On- (top) and Off- event phases (bottom) extracted from (F,G) as heatmap (H) and mean tuning (I). All ROIs are sorted by the timing of the On-event in response to 584 nm (“peak” orange/red) light stimulation. In all heatmaps showing ROIs, lighter colours indicate a higher signal. **J-L**, Activation of the On- (J), Off- (K) and On-Off Contrast-filters (L) for each of 30,000 individual natural spectra (from n = 30 scenes [S6]) plotted against their “brightness”, here computed as their loading against the first principal component (PC) that emerges from PCA across the entire dataset (see also [S6]). Data from individual scenes is indicated by their different coloration. Spearman correlation coefficients ρ as indicated.

## Acknowledgements

Funding was provided by the European Research Council (ERC-StG “NeuroVisEco” 677687), The Wellcome Trust (Investigator Award in Science to TB), The UKRI (BBSRC, BB/R014817/1), the Leverhulme Trust (PLP-2017-005) and the Lister Institute for Preventive Medicine. The authors would also like to acknowledge support from the FENS-Kavli Network of Excellence and the EMBO YIP.

## Author contributions

PB and TB designed the study, with input from FKJ and DO. PB performed 2-photon data collection, pre-processing, and analysis. PB built the light-stimulator with input from FKJ, who also built the volumetric mesoscale two-photon system. PB performed natural imaging data analysis, with input from TB and DO. TB wrote the manuscript with inputs from all authors.

## Supplemental Information

Supplemental Information includes experimental procedures, one figure and one video and can be found with this article online at ….

## Supplemental Information

Document S1. Experimental Procedures and One Figure

Video S1. **Two-photon curved triplane mesoscale imaging of the larval zebrafish brain during hyperspectral full-field stimulation | related to Figure 1A and Figure S1A-C.** Combined responses of one zebrafish’s brain to flashes of different wavelengths of light presented in sequence (cf. Figure 1B, Figure S1D) based on three consecutive scans with three planes each (cf. Figure S1A). Data is averaged over 4 stimulus loops and sped up to 5x real time. In the second video segment, the central panel from the first segment is isolated and montaged to display responses to all 13 tested wavelengths in synchrony, as indicated.

## SUPPLEMENTAL INFORMATION

### SUPPLEMENTAL EXPERIMENTAL PROCEDURES

#### RESOURCE AVAILABILITY

##### Lead Contact

Further information and requests for resources and reagents should be directed to and will be fulfilled by the Lead Contact, Tom Baden.

##### Data and Code Availability

Pre-processed functional 2-photon imaging data, natural imaging data and associated summary statistics, as well as relevant Jupyter notebooks will be made freely available via the relevant links on http://www.badenlab.org/resources and http://www.retinal-functomics.net. The natural imaging dataset was published previously as part of [8].

#### EXPERIMENTAL MODEL AND SUBJECT DETAILS

##### Animals

All procedures were performed in accordance with the UK Animals (Scientific Procedures) act 1986 and approved by the animal welfare committee of the University of Sussex. For all experiments, we used 6-7 *days post fertilization* (*dpf*) zebrafish (Danio rerio) larvae. The following previously published transgenic line was used: Tg(elavl3:H2B-GCaMP6f); ZFIN ZDB-ALT-150916-4 [S1]. Animals were housed under a standard 14:10 day/night rhythm and fed three times a day. For 2-photon *in-vivo* imaging, zebrafish larvae were immobilised in 2% low melting point agarose (Fisher Scientific, BP1360-100), placed on a glass coverslip and submerged in fish water.

##### Light Stimulation

With fish mounted upright, light stimulation was delivered as wide-field flashes from a spectrally broad liquid waveguide with a low NA (0.59, 77555 Newport), positioned next to the objective at ~45°. The other end of the waveguide collected light from 13 “spectrally narrowed” LEDs, as described in detail elsewhere [S2]. All stimuli were series of single LED flashes of light lasting 3 s, separated by gaps of 3 s (1 stimulus loop: 13 LEDs * (3+3) s = 78 s. 3-4 loops were presented and averaged for each recording.

##### 2-photon calcium imaging

All 2-photon imaging was performed on a MOM-type 2-photon microscope (designed by W. Denk, MPI, Martinsried; purchased through Sutter Instruments/Science Products) equipped with a mode-locked Ti:Sapphire laser (Chameleon Vision-S, Coherent) tuned to 960 nm for SyGCaMP imaging. We used one fluorescence detection channel (F48×573, AHF/Chroma), and a water immersion objective (W Plan-Apochromat 20x/1,0 DIC M27, Zeiss). For image acquisition, we used custom-written software (ScanM, by M. Mueller, MPI, Martinsried and T. Euler, CIN, Tuebingen) running under IGOR pro 6.3 for Windows (Wavemetrics).

To expand the field of view to ~1.2 mm diameter, which allowed capturing the entire brain’s length in a single scan, we used a non-telecentric optical approach as described in detail elsewhere [S3]. The excitation spot (point spread function) in this configuration was ~0.7 μm (xy) and ~11 μm (z) at full width half maximum. This optical configuration can in principle capture the signals from individual larval zebrafish somata [S3]. However, in this work it was our intention to capture the bulk spectral responses across large fractions of the brain. Accordingly, we balanced recording area and spatial sampling such that individual somata effectively corresponded to single, or at most groups of 2-4 pixels (3 planes covering ~450×1,000 μm with a 160×350 px scan each to yield ~2.9 μm voxel xy-spacing, compared to average zebrafish neuronal soma diameter of ~7 μm; 1 ms per line, 2.08 Hz volume rate).

To follow the brain’s natural 3D curvature, we also systematically 3D-bent each scan-plane as a function of the slow scanning-mirror’s position to form a “half-pipe”. Curvature was achieved via rapid remote focussing synchronised with the scan pattern, as described in detail elsewhere [S3]. The degree of peak axial curvature was empirically adjusted between 0-150 μm between scans and planes to achieve best overall sampling of the entire brain.

##### Pre-processing and extraction of response amplitudes of 2-photon data

Recordings were linearly interpolated to 42 Hz and manually aligned between fish using a time-averaged brightness projection. Regions of interest (ROIs), corresponding to individual and/or small groups of neighbouring neuronal somata were defined automatically using custom Python scripts. In short, we used a “quality-index” (QI, described in detail elsewhere [S4]) to first identify individual pixels that exhibited reliable responses to repeated stimulation. For this, we computed a pixel-wise QI-projection of the deinterleaved recording, sorting QI-pixels in descending order. The resulting curve was differentiated using *scipy.interpolate.splrep*. Pixel indices between inflections of the differential were projected back into space. Contours were identified using dilation (3,3)-erosion(2,2) and contour finding of Python-OpenCV. Individual contours were taken as ROIs, discarding any ROIs with a diameter > 15 μm. QI per ROI was then recalculated and used for further thresholding at QI>0.5. From here, fluorescence traces were extracted and z-normalized based on the 6 s at the beginning of recording prior to stimulus presentation. Overall, this strategy served to balance the need to combine multiple pixels into ROIs to boost their signal-to-noise, with a goal of keeping ROIs as small and localised as possible to approximately report the signals single, or from at most very small groups of somata that responded in a similar manner. This compromise was necessary to accommodate the large size of the scan pattern capturing the entire length of the brain while also maintaining a reasonable imaging rate. A stimulus time marker embedded in the recording data served to align the traces relative to the visual stimulus with a temporal precision of 1 ms.

##### Separation of On- and Off responses

Calcium traces were deconvolved using ARMA(1) (caiman.source_extraction.cnmf.deconvolution, [S5]). Inferred discrete events were partitioned into events occurring during stimulus presentation and the complement.

##### Computing the brain’s bulk spectral tuning functions

Inferred events were summed over respective stimulus time windows. Sums were averaged over all recorded traces. Contrast between On and Off portions of the response was calculated as their difference over their sum.

##### Natural Imaging Data Analysis

Hyperspectral data were obtained from [S6] and element-wise multiplied with a deuterium light source derived correction curve [S.x]. The data were restricted to the domain of 360-650 nm. Here, the long-wavelength end of the domain was decided based on the long-wavelength opsin absorption curve; the short-wavelength end was dictated by the sensitivity of the spectrometer. Spectra were scaled by standard deviation within a given scene. Traces were multiplied with the respective On- and Off-filters. The responses were summed within spectrum to produce a single number per point spectrum (or 800-long vector per scan). These vectors were standard-deviation-scaled within a scene. Spatial projections of filter responses were Gaussian-smoothed in space (σ=2px).

## Notes

### Competing Interest Statement

The authors have declared no competing interest.

